# Structural basis for reduced ribosomal A-site fidelity in response to P-site codon-anticodon mismatches

**DOI:** 10.1101/2023.01.28.526049

**Authors:** Ha An Nguyen, Eric D. Hoffer, Crystal E. Fagan, Tatsuya Maehigashi, Christine M. Dunham

**Author notes:** Contact: Christine M. Dunham. **Data deposition:** X-ray crystallography, atomic coordinates, and structure factors have been deposited in the Protein Data Bank, www.pdb.org (PDB codes 8FOM, 8FON).

## Abstract

Rapid and accurate translation is essential in all organisms to produce properly folded and functional proteins. mRNA codons that define the protein coding sequences are decoded by tRNAs on the ribosome in the aminoacyl (A) binding site. The mRNA codon and the tRNA anticodon interaction is extensively monitored by the ribosome to ensure accuracy in tRNA selection. While other polymerases that synthesize DNA and RNA can correct for misincorporations, the ribosome is unable to correct mistakes. Instead, when a misincorporation occurs, the mismatched tRNA-mRNA pair moves to the peptidyl (P) site and from this location, causes a reduction in the fidelity at the A site, triggering post-peptidyl transfer quality control. This reduced fidelity allows for additional incorrect tRNAs to be accepted and for release factor 2 (RF2) to recognize sense codons, leading to hydrolysis of the aberrant peptide. Here, we present crystal structures of the ribosome containing a tRNA^Lys^ in the P site with a U•U mismatch with the mRNA codon. We find that when the mismatch occurs in the second position of the P-site codon-anticodon interaction, the first nucleotide of the A-site codon flips from the mRNA path to engage highly conserved 16S rRNA nucleotide A1493 in the decoding center. We propose that this mRNA nucleotide mispositioning leads to reduced fidelity at the A site. Further, this state may provide an opportunity for RF2 to initiate premature termination before erroneous nascent chains disrupt the cellular proteome.

## INTRODUCTION

The accurate flow of genetic information is vital for cellular life. DNA and RNA polymerases copy nucleic acid templates into complementary nucleic acids and Watson-Crick base pairing between these nucleotide strands guides accuracy. The thermodynamic differences between base pairings alone cannot fully account for the exceptional accuracy of replication (∼10^−9^) and transcription (∼10^−5^) (1). To accomplish high accuracy, both DNA and RNA polymerases detect misincorporations by proofreading mechanisms that excise the incorrect nucleotide and replace with the correct nucleotide. Continuous replication or transcription is then maintained without having to discard the current product and restart. In contrast, during protein synthesis, the incorporation of an incorrect amino acid is irreversible. This irreversibility arises in part because the template (mRNA codon) and product (amino acids) are different chemical moieties preventing the use of Watson-Crick base pairing as a mechanism to retroactively determine if the product is correct. An additional challenge is that the large distance (∼70 Å) between the mRNA-tRNA base pairing in the decoding center on the small ribosomal subunit and the peptidyl transferase center (PTC) on the large ribosomal subunit where aminoacyl groups attached to tRNAs are added to the nascent chain, prevents a rapid response (2,3). Collectively, these differences may account for higher error rates in protein synthesis (4).

The ribosome maintains sufficient fidelity during protein synthesis for production of a functional proteome. Using both kinetic proofreading and induced fit mechanisms, the ribosome rapidly selects the correct tRNA substrate from incorrect but structurally similar tRNAs (5-9). Ternary complexes containing aminoacyl-tRNAs (aa-tRNAs), EF-Tu and GTP (aa-tRNA•EF-Tu•GTP) are delivered to the aminoacyl (A) site of the ribosome and encounter two kinetic checkpoints before acceptance. First, Watson-Crick base pairing between the codon and anticodon is inspected during a process called initial codon selection. The ribosomal A site has an extensive monitoring network comprised of ribosomal 16S RNA (rRNA) nucleotides G530, C1054, A1492, A1493 and A1913 (*E. coli* numbering) which undergo conformational changes to directly inspect the pairing of the codon-anticodon on the 30S subunit (Fig. 1) (10-12). The first two positions of the codon-anticodon interaction are required to be Watson-Crick (A-U or G-C) due to the constraints of the A site, while the third position can either be Watson-Crick or an interaction that resembles the geometry of a Watson-Crick pairing (e.g., G•U pairing or the pairing of a modified anticodon nucleotide with an mRNA nucleotide). The complementarity of the codon-anticodon interaction stabilizes the ternary complex while incorrect aa-tRNAs rapidly dissociate (13,14). Second, a correct Watson-Crick base pair causes rapid hydrolysis of GTP by EF-Tu while incorrect pairings induce slower GTP hydrolysis and EF-Tu disassociation (15). Rapid hydrolysis also enables conformational changes in the ternary complex leading to full accommodation of aa-tRNAs on the large 50S subunit. These steps ensure high accuracy and speed during protein synthesis.

**Figure 1.**
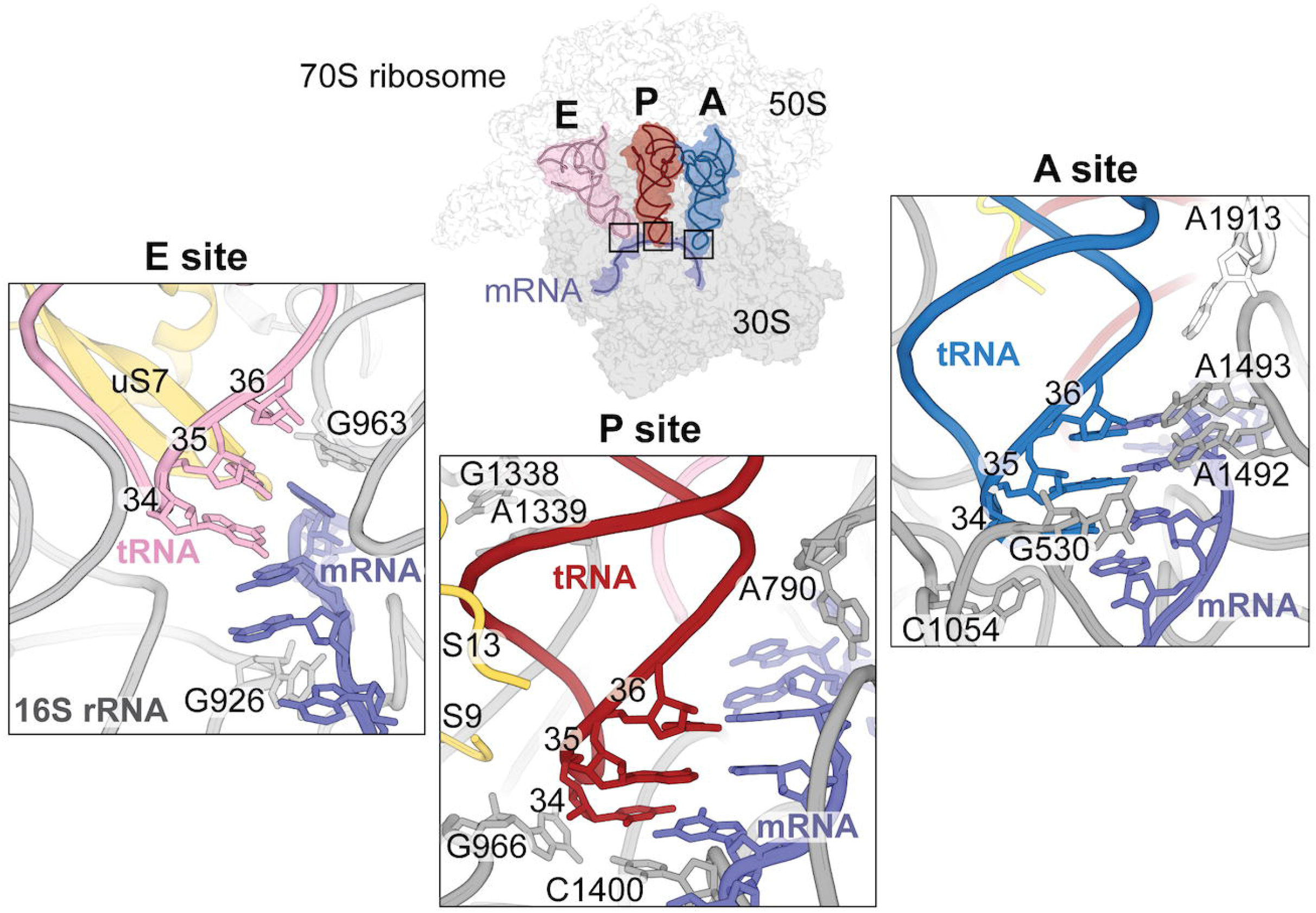
The ribosome extensively probes the tRNA-mRNA interaction at the A site but does not monitor P- and E-site mRNA-tRNA base pairing. tRNA binding sites on the 70S ribosome are shown (PDB code 6OF6). tRNAs enter the ribosome first at the A site (right panel) and 16S rRNA nucleotides A1492, A1493, C1054 and G530 (gray), and 23S rRNA A1913 nucleotides monitor the mRNA (lavender) - tRNA (blue) pairing. After peptidyl transfer and movement to the P site (middle panel), the P-site tRNA (red) is gripped by two 16S rRNA G1338-A1339 and A790, and the C-terminal tails of proteins uS9 and uS13 (yellow). G966 and C1400 surround nucleotide 34 of the tRNA. In the E site (left panel), the ribosome makes minimal contacts with the tRNA (pink) and mRNA (lavender). Ribosomal protein uS7 (yellow) contacts the tRNA and 16S rRNA nucleotides G963 and G926 interact with the mRNA.

Despite proofreading mechanisms, missense errors still occur *in vivo* at a rate of one in ∼3000 amino acids incorporated (16,17), which is notably lower than the *in vitro* error rate of 1 in 500 amino acids (1). This discrepancy implies additional quality control processes exist beyond A-site tRNA selection. Indeed, the discovery of a post-<u>p</u>eptidyl transfer <u>q</u>uality <u>c</u>ontrol mechanism (post-PT QC) revealed that codon-anticodon mismatches that bypass A-site surveillance mechanisms influence the next round of tRNA selection from their position in the peptidyl (P) site (18). Using a well-known *in vivo* misincorporation event where tRNA^Lys^ miscodes at high levels on the near-cognate asparagine AAU codon (19,20), when the mismatched codon-anticodon pair moves to the P site, a subsequent loss in fidelity at the A site ensues (a near-cognate interaction between the codon and anticodon is defined as two Watson-Crick base pairs and a single non-Watson-Crick base pair) (18,21,22). This loss of fidelity at the A site causes tRNA selection errors followed by premature termination mediated by release factors 2 (RF2) recognition of non-stop codons (1). Incorrect tRNA selection at the A site in this context causes an accelerated rate of GTP hydrolysis by EF-Tu with tRNA accommodation occurring at similar rates to those of correct tRNAs (21). Premature termination by RF2 on non-stop codons is two orders of magnitude higher after a single misincorporation event and four orders of magnitude higher after two consecutive misincorporation events (1,23). The post-PT QC response is influenced by the identity of the mismatched codon-anticodon pairs suggesting that the ribosome can discriminate between subtle differences in mismatches (18). For example, first or second codon position G•U and U•U mismatches robustly activate post-PT QC however, not all mismatches at the third codon-anticodon position elicit post-PT QC. For example, while a third-position U•U mismatch activates post-PT QC, a G•U mismatch does not trigger premature termination (18). The extent of post-PT QC is also dependent on the position of the mismatch, with RF2-mediated peptidyl hydrolysis on non-stop codons the greatest with second codon-anticodon position mismatches in the P site (1,18,21).

The codon-anticodon pairing in the ribosomal P site is not stringently monitored for Watson-Crick base complementarity unlike in the A site where tRNAs are selected (Fig. 1). Instead, the P site optimally recognizes initiator tRNA^fMet^ to begin protein synthesis and during the elongation phase of translation, the ribosome “grips” elongator tRNAs at the anticodon stem to aid in ensuring the frame of the mRNA is maintained. 16S and 23S rRNA nucleotides and ribosomal proteins form a network of interactions that surround the P-site tRNA but there is minimal direct inspection of the anticodon-codon interaction (24). 16S rRNA nucleotides G966 and C1400 form a non-Watson-Crick interaction that packs beneath the third position of the codon-anticodon while G1338 and A1339 surround anticodon stem G-C base pairs to distinguish between initiator and elongator tRNAs and prevent premature tRNA translocation from the P to the exit (E) site (25,26). C-terminal tails of ribosomal proteins uS9 and uS13 also contact the tRNA anticodon stem loop (ASL) (Fig. 1) (“u” refers to a universal ribosomal protein found in all three kingdoms (27)). These interactions collectively stabilize already selected tRNAs, but how the ribosome recognizes P-site codon-anticodon mismatches is unclear given these minimal interactions with the codon-anticodon pair.

To determine how the ribosome recognizes mismatches in the P site, we solved two structures of 70S ribosomes bound to different mRNAs containing a single U•U mismatch at either the first or second codon-anticodon position (Fig. 2). These ribosome structures contain tRNA^Lys^ (anticodon is SUU, where S is 5-methylaminomethyl-2-thiouridine (mnm5s2U)) bound to either a mismatched UAA or AUA codon in the P site along with a cognate tRNA-mRNA pair in the A site (Fig. 1). The pairing of tRNA^Lys^ with both these codons causes high levels of miscoding leading to post-PT QC (18,20,21). In both cases, the U•U mismatches in the codon-anticodon interaction still forming base paired stacks even though the distances are too wide to allow for Watson-Crick pairing. In the structure containing a second U•U mismatch, the first nucleotide of the A-site mRNA codon flips ∼90° away from the mRNA path towards A1493 of the decoding center. This movement leaves the A-site mRNA codon with only two of the three nucleotides properly positioned to interact with either incoming tRNAs or RFs. In contrast, the first position codon-anticodon mismatch in the P site does not influence the position of the A-site codon, consistent with biochemical studies that demonstrate second position mismatches trigger post-PT QC at higher levels than mismatches at the first or the third position (1,18). We propose that the mispositioning of the A-site codon induced by the P-site mismatch serves as a signal to trigger premature termination and post-PT QC to prevent further erroneous protein synthesis.

**Figure 2.**
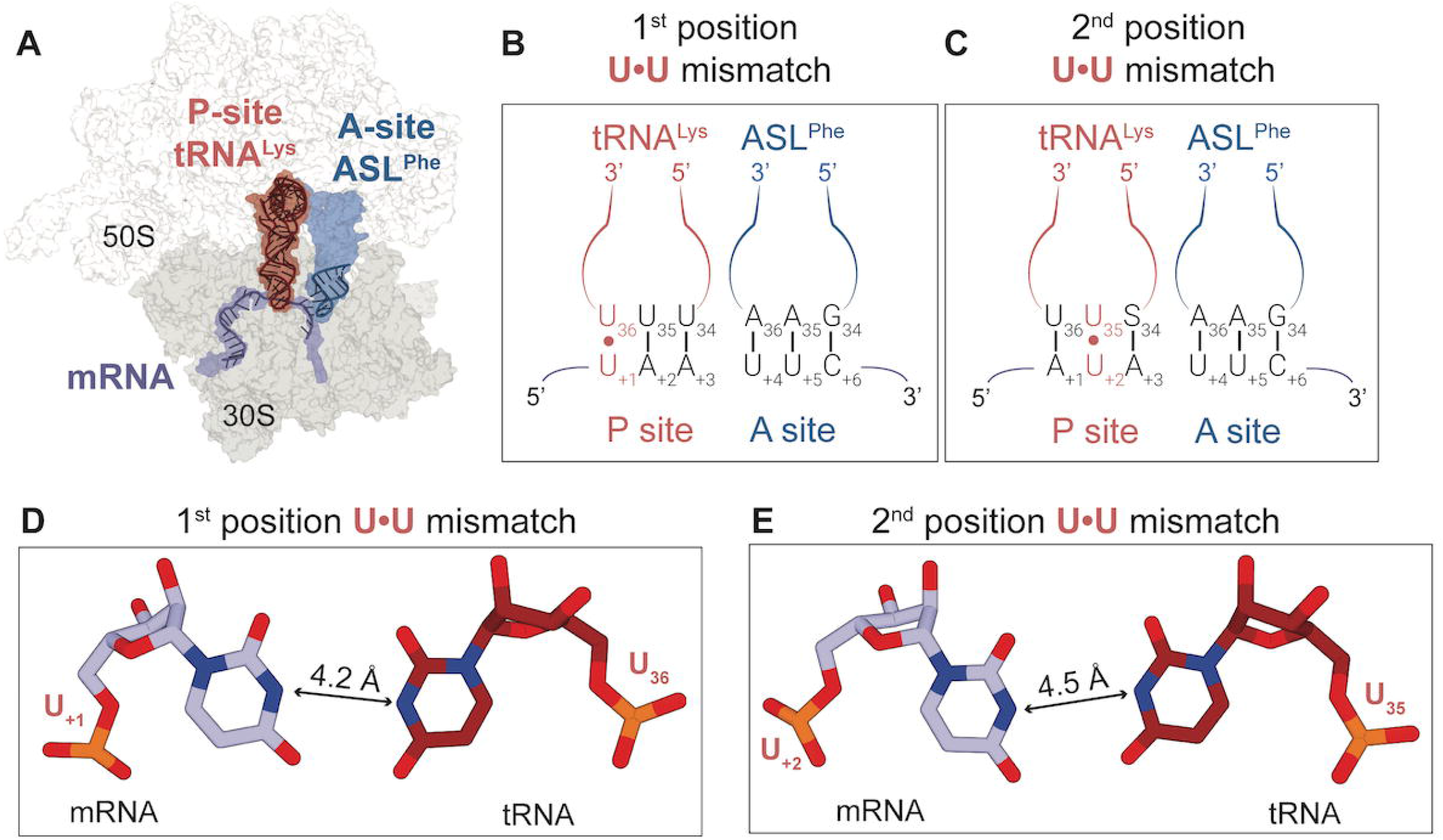
Structures of the 70S ribosome containing codon-anticodon mismatches in the peptidyl (P) site. A. An overview of the 70S ribosome complexes in this study showing the mRNA (lavender) along with the tRNA^Lys^ (red) in the P site and the anticodon stem loop (ASL, blue) of tRNA^Phe^ in the A site. The full binding sites for the mRNA and tRNAs are shown in surface in their corresponding colors. B. tRNA^Lys^ binds to the UAA codon resulting in a 1^st^ position U•U mismatch in the P site. Nucleotide numbering for the mRNA starts at +1 as the first position of the P site codon. C. tRNA^Lys^ binds to the AUA codon resulting in a 2^nd^ position U•U mismatch in the P site. S34 in tRNA^Lys^ refers to 5-methylaminomethyl-2-thiouridine (mnm^5^s^2^U). D. The 1^st^ position U•U mismatch is Watson-Crick-like in its orientation but is too wide for hydrogen bonding. E. Similar to the 1^st^ position mismatch, the 2^nd^ position U•U mismatch also adopts a wider Watson-Crick-like base pairing.

## RESULTS

### Single codon-anticodon nucleotide mismatches minimally impact the architecture of the P site

To understand how P-site mismatches between the mRNA and tRNA influence fidelity at the A site, we solved two crystal structures of *Thermus thermophilus* ribosomes containing tRNA^Lys^ bound to either a UAA and AUA codon, creating a U•U mismatch at either the first or second positions of the codon-anticodon interaction (Fig. 2A-C). In both structures, we find no apparent differences in the overall P-site architecture compared to ribosome structures containing a cognate mRNA-tRNA pair: 16S rRNA nucleotides G966 and C1400 pack beneath the third position of the codon-anticodon, G1338 and G1339 form A-minor interactions with the anticodon stem nucleotides G30-C40, G29-C41 of the tRNA, and uS9 and uS13 tails extend into the P site (Fig. S1). In both structures, the U•U mismatch between the P-site tRNA^Lys^ and the mRNA codon is either at the first position of the UAA codon (U_+1_•U_36_) or second position of the AUA codon (U_+2_•U_35_) (+1 numbering starts at the first mRNA nucleotide in the P site codon; U_36_ and U_35_ refer to the anticodon positions) (Figs. S2, S3). Similar to previous structures of G•U codon-anticodon mismatches in the P site (28), both U•U pairs mimic the geometry of a Watson-Crick base pair even though the nucleobases are too far apart for hydrogen bonding (Figs. 2D-E). However, this geometry allows the U•U mismatches to still form stacking interactions within the codon-anticodon. Thus, in contrast to the various structural reorganizations that can occur in the A site when a codon-anticodon mismatch is present at the first or second codon-anticodon position (11,29,30), the P site is unaltered despite the codon-anticodon mismatches (Fig. S1).

### Second nucleotide U_+2_•U_35_ mismatch in the P-site codon-anticodon causes the first U_+4_ nucleotide of the A-site codon to deviate from the mRNA path

In both 70S structures containing a P-site U•U mismatch at the first (U_+1_•U_36_) or second (U_+2_•U_35_) position of the codon-anticodon, the A site contains an ASL^Phe^ bound to a cognate UUC phenylalanine codon along with CC-puromycin, a 3’-CCA end mimic of tRNA that binds to the 50S A site and prevents other tRNAs binding the A site. Once a codon-anticodon mismatch is present in the P site, CC-puromycin was required to block full-length tRNA^Lys^ nonspecifically binding at the A site. In the 70S structure containing a P-site U_+1_•U_36_ mismatch at the first position, the interaction between the codon-anticodon in the A site contains three standard Watson-Crick base pairs (A_36_-U_+4,_ A_35_-U_+5_, G_34_-C_+6_) (Fig. 3B). In this context, 16S rRNA decoding center nucleotides A1492, A1493, and G530 interact with the first and second codon-anticodon base pairs directly to probe for Watson-Crick pairs, and C1054 stacks with the third base pair. In the 70S structure containing a P-site U_+2_•U_35_ mismatch at the second position, the first nucleotide of the mRNA codon in the A site (U_+4_) deviates from its normal position and appears to flip ∼90° from the mRNA path (Figs. 3C, 4A and Fig. S4). Normally U_+4_ would form a base pair with A_36_ of the A-site tRNA as the first base pair of the A-site codon-anticodon interaction. While the ribose of U_+4_ fits the experimental 2f_o_-f_c_ density well, there is minimal density for the nucleobase. We tried modeling the A_36_-U_+4_ interaction as Watson-Crick but there was significant difference density surrounding the nucleobase, strongly suggesting the model was incorrect (Fig. 3D and Fig. S4). In contrast, there is little to no difference density surrounding U_+4_ when modeled flipped away from the mRNA path (Fig. 3D). We therefore modeled U_+4_ as flipped where it would make direct interactions with 16S nucleotide A1493 of the decoding center (Fig. 4B). Specifically, the 2’-OH of U_+4_ hydrogen bonds with the 2’-OH and the phosphate oxygen of A1493. We acknowledge that U_+4_ may be flexible which does not usually occur because of the strong tendency of the tRNA to pair with the mRNA.

**Figure 3.**
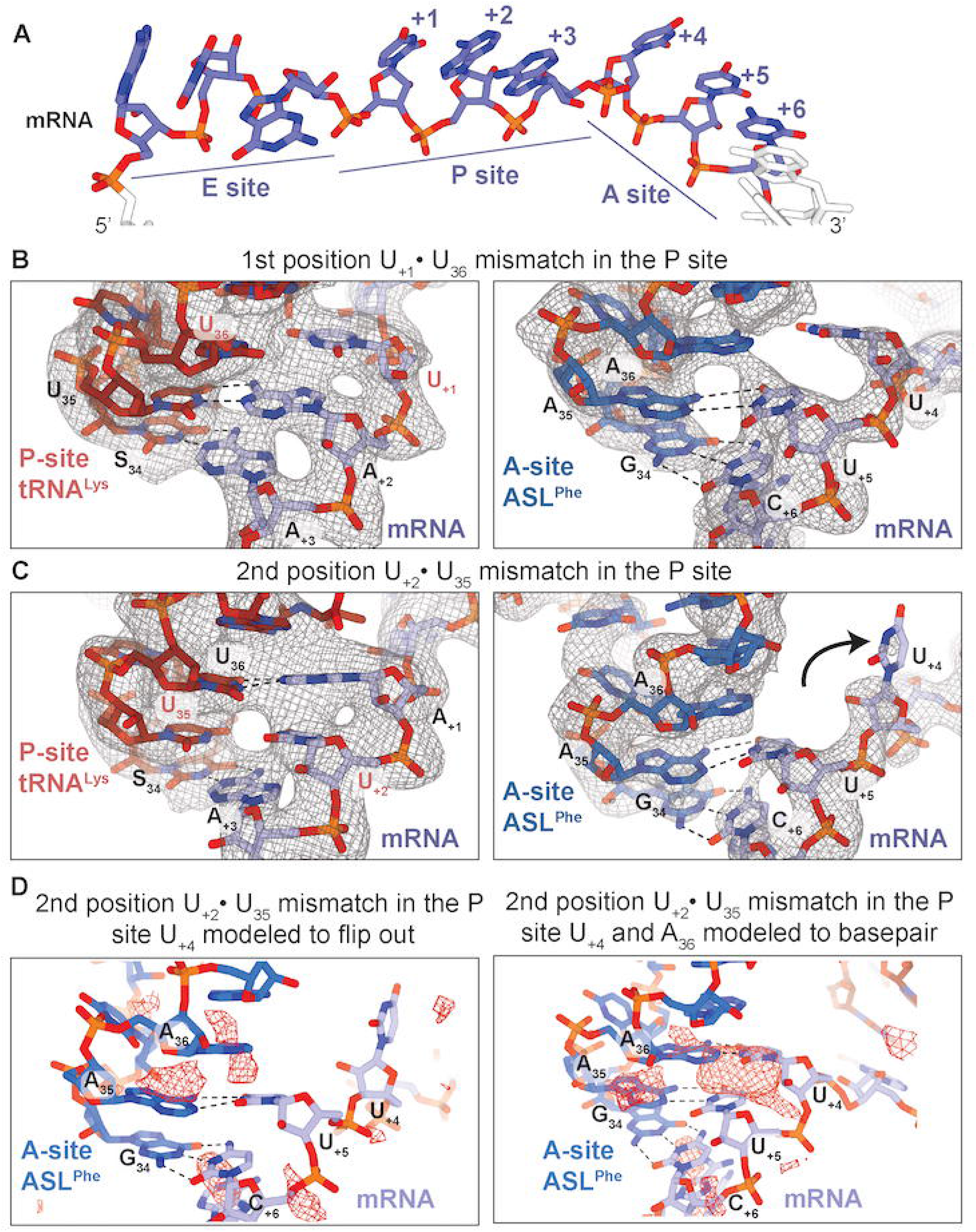
The context of the P-site mismatch results in mRNA mispositioning. A. The typical path of the mRNA through the ribosome showing the codon positioning in the ribosomal tRNA binding sites: exit (E) site, peptidyl (P) site, aminoacyl (A) site. The nucleotides are numbered with the first nucleotide in the P site denoted +1. B. When the U•U mismatch is at the first nucleotide of the P-site codon-anticodon pairing, the mRNA (lavender) is positioned correctly in both the P and A sites. The P-site codon nucleotides (U_+1_, A_+2_, A_+3_) is paired to the tRNA^Lys^ (red) anticodon (S_34_U_35_U_36_) (left). The A-site codon nucleotides form three Watson-Crick base pairs to the A-site ASL^Phe^ (blue) anticodon (U_+4_-A_36_, U_+5_-A_35_, C_+6_-G_34_) (right). C. When the U•U mismatch is located at the second position of the P-site mRNA-tRNA codon-anticodon interaction, the mRNA is mispositioned in the A site. The P-site codon (U_+1_, A_+2_, A_+3_) is correctly paired to the P-site tRNA^Lys^ (left). The first nucleotide of the A-site codon (U_+4_) is flipped away from the A-site ASL^Phe^ causing the A-site anticodon-codon interaction is disrupted (right). Only two Watson-Crick base pairing interactions are maintained: U_+5_-A_35_, C_+6_-G_34_. 2F_o_-F_c_ electron density maps contoured at 1.0σ are shown in gray mesh. D. Modeling of the U_+4_-A_36_ interaction as non Watson-Crick and with U_+4_ flipped from the mRNA path shows little to no F_o_-F_c_ difference electron density (left; red mesh). Modeling the U_+4_-A_36_ interaction as Watson-Crick reveals significant F_o_-F_c_ difference electron density indicating the nucleobase likely does not occupy this position (right). F_o_-F_c_ electron density maps are contoured at 2.5σ.

**Figure 4.**
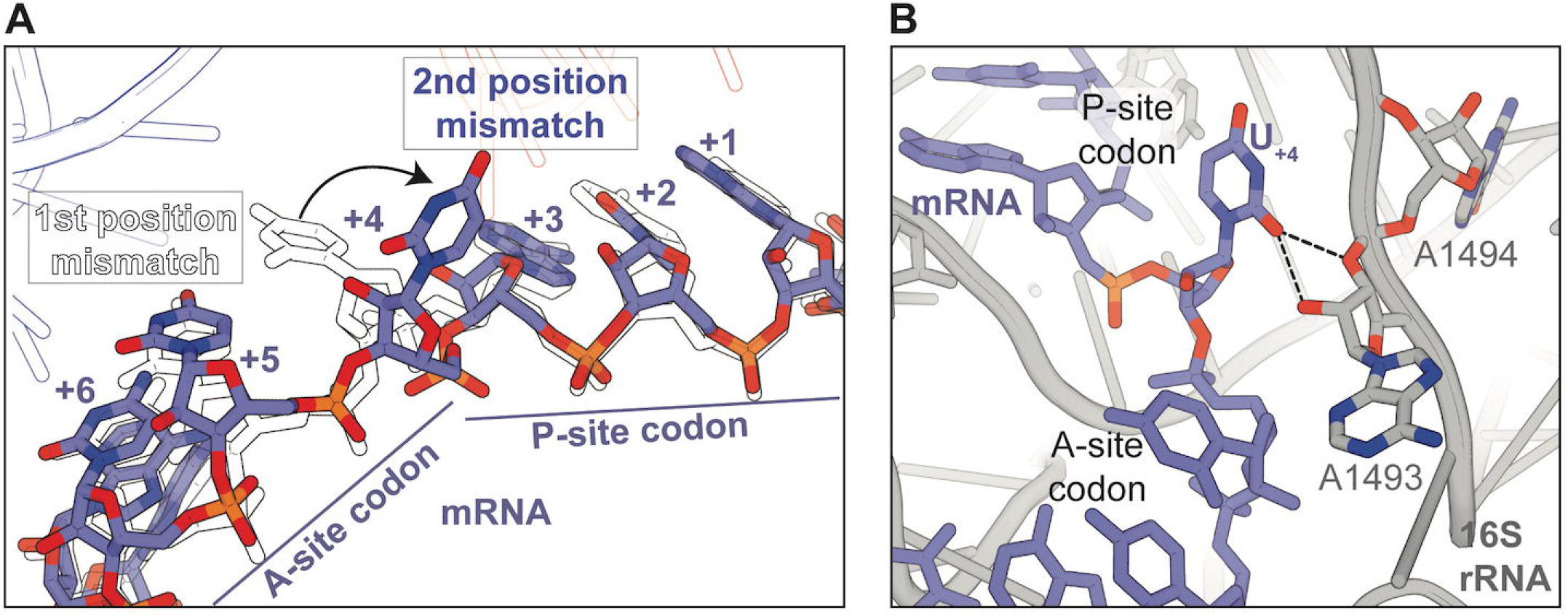
The mRNA mispositioning disrupts the A-site codon-anticodon interactions. A. Overlay of the mRNA paths from both the structures indicate the backbone paths are very similar. The first nucleotide of the A site is flipped away by ∼90° from the normal position when the mismatch is at the second position of the P-site anticodon-codon interaction. The nucleotides from the 1^st^ position mismatch are in black outlines, and the solid nucleotides represent the 2^nd^ position mismatch structure. B. When there is a 2^nd^ position U•U mismatch in the P site, the first nucleotide of the A site (U_+4_) flips and interacts with 16S rRNA A1493 instead of pairing with the tRNA anticodon.

## DISCUSSION

In this study, we examined the structural basis for how P-site U•U mismatches between the codon and anticodon affect the fidelity at the adjacent A site. It was previously shown that when an incorrect codon-anticodon pairing escapes rejection at the A site, its incorrect aminoacyl group is added to the growing nascent chain, the mismatched mRNA-tRNA pair is translocated to the P site, and quality control mechanisms are initiated to reduce fidelity at the A site (18,21). This remarkable reduction in A-site fidelity allows for incorrect tRNAs to be accepted by the ribosome and causes subsequent premature termination from RF2 recognition of non-stop codons. Hydrolysis of this erroneous polypeptide is necessary to stop the synthesis of aberrant polypeptides and allow the ribosome to be recycled for further rounds of translation.

Our structures here investigate two particular instances where tRNA^Lys^ miscodes at high levels on either the UAA ochre stop codon (first position U•U mismatch) or the AUA isoleucine codon (second position U•U mismatch) at rates of 4.1 and 3.5 × 10^−4^, respectively *in vivo* (20). Once this miscoding happens at the A site, the mRNA-tRNA mismatched pair is translocated to the P site where this mismatch still can influence the fidelity of decoding at the A site. Both these first and second U•U mismatches cause reductions in A-site fidelity allowing high levels of termination on non-stop codons (18). However, the rate of hydrolysis is ∼3 fold higher when the mismatch is in the second position of the P site as compared to the first position (18), consistent with the structural changes we observe. Also consistent with this, the selection efficiency of the ribosome to detect single mismatches during miscoding by tRNA^Lys^ in the A site has the highest discrimination when U•U mismatches are present at the second position, followed by the first and third position mismatches (31). This critical importance of second-position translational accuracy suggests a potential explanation for the mRNA flipped nucleptide we observe in the structure with the more impactful second position U•U mismatch but not for the structure with the U•U mismatch in the first position.

The movement of U_+4_ from the normal mRNA path in the A site is surprising for several reasons. First, the A-site codon-anticodon interaction is cognate and three Watson-Crick base pairs should form. Second, when there is a U_+1_•U_36_ mismatch in the first position of the P site, the mRNA path in the A site is unaffected (Figs. 2B,3B). However, these structures are consistent with the biochemical characterization of the post-PT QC response that reveals a more robust response to second position mismatches (18). We propose that the second position U_+2_•U_35_ mismatch causes a reduction in fidelity through the mispositioning of the first nucleotide of the A-site codon. The impaired mRNA presentation in the A site may allow binding of non-cognate tRNAs and release factors and ultimately induce premature termination.

Prior ribosome structures of single P-site G•U mismatches at all three positions of the codon-anticodon show these pairs adopt Watson-Crick-like geometry (32). These structures initially suggested that few differences exist between how A or P site mismatches are recognized by the ribosome. However, biochemical studies strongly suggest that these differences could be tRNA^Lys^ dependent, which is known to miscode at high levels and seems to be the ideal tRNA to understand this phenomenon (18-20). One difference in the U•U mismatch structures presented here and other G•U mismatch structures is the distance between the mismatched codon and anticodon. The G•U mismatches form hydrogen bonding interactions but the distance is too great between U•U mismatches to form electrostatic interactions.

Many miscoding studies have focused on tRNA^Lys^ because of its well-known propensity to miscode on codons containing a mismatch at the third codon-anticodon position such as the asparagine AAU codon (19,20). Quantifying every possible combination of codon mismatches that cause misreading by tRNA^Lys^ reveals no clear trends; both the mismatch position and the type of nucleotide mismatches themselves do not appear to influence miscoding. Instead, the codon family seems to be the main determinant for high levels of miscoding by tRNA^Lys^ (20). For example, tRNA^Lys^ has a high misreading rate on arginine (AGA, second position G_+2_•U_35_ mismatch; AGG, second position G_+2_•U_35_ and third position G_+3_•U_34_ mismatches), asparagine (AAU, third position U_+3_•S_34_), and termination (UAG, first position U_+1_•U_36_ and third position G_+3_•U_34_ mismatches) codons. Expansion of the work to other tRNAs indicates that miscoding rates are highest for U•U mismatches (33). Since the ribosomal A site heavily constrains the formation of the codon-anticodon interaction, perhaps the smaller pyrimidine-pyrimidine mismatch in U•U mismatches bypasses A-site proofreading mechanisms more easily because of a lack of steric clashes or unfavorable backbone torsions present in wider pyrimidine-purine or purine-purine mismatches (34). In both P-site U•U mismatch structures presented here, the U•U pairings are wider as compared to normal Watson-Crick pairings (>4 Å between uracil Watson-Crick edges vs ∼3.2 Å for a canonical A-U base pair). Despite this widening, the nucleobases of the U•U mismatch are positioned for Watson-Crick-like pairing (Figs. 2D-E). These U•U mismatches also do not cause major perturbations in the shape of the anticodon stem loop suggesting their influence appears to be localized to the codon and anticodon (Figs. S1-3).

Protein synthesis terminates when the ribosome reaches a stop codon in the mRNA reading frame and RFs hydrolyze the nascent chain from the tRNA (Fig. 5). Bacteria contain two RFs (RF1 and RF2) that have different and overlapping stop codon recognition: RF1 recognizes UAA and UAG and RF2 recognizes UAA and UGA. The sense or stop codon mRNA in the A site adopts a different orientation dependent upon RF binding. When tRNAs bind, three planar base pairs form a π -stacking interaction. RF binding causes stacking of only the first two positions of the codon while the third nucleotide flips and stacks with 16S rRNA. The decoding center also does not participate in RF recognition of stop codons and instead stabilizes the catalytically active ‘open’ conformation of RFs.

**Figure 5.**
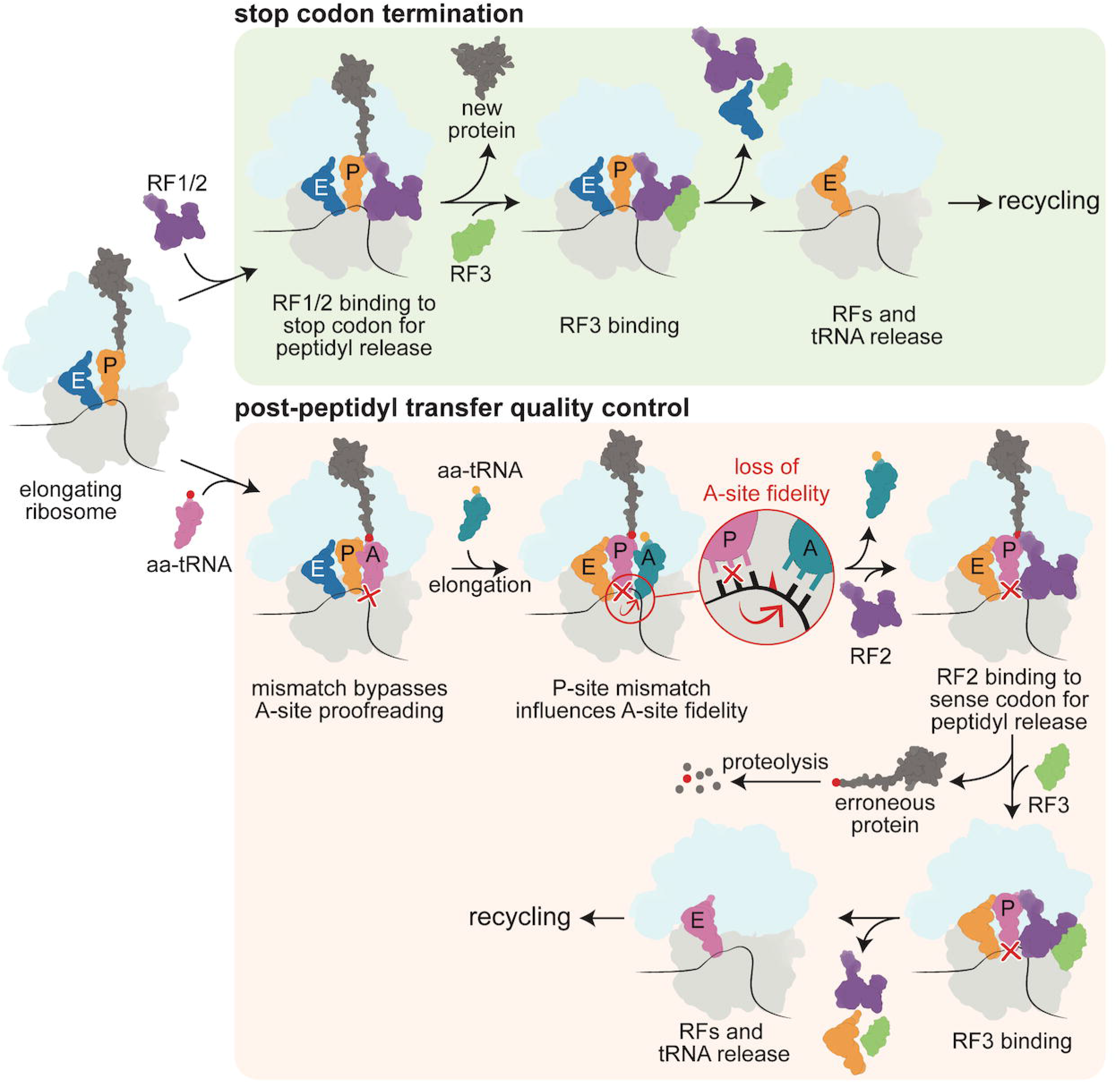
Mismatches in the P site trigger post-peptidyl transfer quality control. During canonical translation termination (top pathway), release factors RF1 or RF2 (pink) recognize when the ribosome reaches a stop codon. They bind to the stop codon in the A site and extends into the peptidyl transferase center to hydrolyze the nascent peptide chain (yellow) still bound to P-site tRNA. RF3 then binds to the ribosome complex to cause RF1/2 to dissociate from the ribosome and allow for the recycling of the entire translation machinery. When the elongating ribosome encounters a mismatch that bypassed A-site proofreading and an incorrect amino acid is incorporated into the nascent chain (yellow), post-peptidyl transfer quality control is triggered (bottom pathway). Mismatches in the P site cause a loss of fidelity in the A site and encourage successive errors to be incorporated. RF2 is then able to recognize these ribosome complexes and cause premature termination on a sense codon. The erroneous protein is quickly degraded, and the ribosome undergoes the rest of the termination pathway to be recycled.

The P-site codon-anticodon mismatch structures presented here contain a cognate interaction in the A site between the phenylalanine UUC codon and tRNA^Phe^. Despite this, three Watson-Crick pairs fail to form when the P-site mismatch is at the second position (U_+2_•U_35_). Interestingly, only RF2 is active in post-PT QC (35) and while RF recognition of stop codons is well understood, this leads to many outstanding questions of how RF2 could mediate premature termination. We would anticipate higher plasticity in how RF2 accommodates changes in the position of the A-site codon and that inclusion of RF3 likely contributes to higher premature termination on sense codons by expanding the recognition capacity of RFs.

## MATERIALS AND METHODS

*T. thermophilus* 70S ribosomes were purified as previously described with slight modifications (36). Ribosome complexes were formed sequentially by adding 4.4 µM 70S ribosomes, 8.8 µM mRNA (IDT), 11 µM CC-puromycin (IDT), 11 µM P-site tRNA^Lys^ (Chemical Block), and 22 µM ASL^Phe^ (IDT) in buffer (5 mM HEPES-KOH pH 7.5, 50 mM KCl, 10 mM NH_4_Cl, 10 mM Mg(CH_3_COO)_2_, 6 mM β-mercaptoethanol) at 55 °C (Table S2). CC-puromycin is an RNA-antibiotic conjugate that mimics the CCA-end of aa-tRNAs that bind normally at the A site. The initial addition of CC-puromycin is necessary to prevent tRNA^Lys^ from binding at the A site and instead positions tRNA^Lys^ exclusively in the P site. Ribosome crystals were grown in 0.1 M Tris-HOAc pH 7.0, 0.2 M KSCN, 4-4.5% (v/v) PEG 20K, 4.5-5.5% (v/v) PEG 550MME, 10 mM Mg(OAc)_2_ and 2.8 µM Deoxy BigCHAP. Crystals were cryoprotected using a stepwise series of cryoprotectants (0.1 M Tris-HOAc pH 7.0, 0.2 M KSCN, 5.5% (v/v) PEG 20K, 10/20/30% (v/v) PEG 550MME, 10 mM Mg(OAc)_2_) before being flash frozen in liquid nitrogen. X-ray diffraction data sets were collected at the 24ID-C NE-CAT beamline at the Advanced Photon Source at Argonne National Laboratory. Datasets were processed in XDS (37) and molecular replacement performed using PDB code 4Y4O as the starting model with PHENIX (38),(39). The coordinates were finalized through iterative rounds of refinements in PHENIX and model building in Coot (40). Figures were created in PyMol (41).

## ACKNOWLEDGMENTS

We thank Dr. Graeme L. Conn and Dunham laboratory members for their critical reading of the manuscript. Support for this work was provided by the Department of Defense through the National Defense Science and Engineering Graduate Fellowship Program (CEF), NIH Training Grant T32 GM8367 (CEF), and NIH R01GM093278 (CMD). This work is based upon research conducted at the Northeastern Collaborative Access Team beamlines, which are funded by the National Institute of General Medical Sciences from the National Institutes of Health (P30 GM124165). This research used resources of the Advanced Photon Source, a U.S. Department of Energy (DOE) Office of Science User Facility operated for the DOE Office of Science by Argonne National Laboratory under Contract No. DE-AC02-06CH11357.

