## Supplemental Data File for "Structural basis for reduced ribosomal A-site fidelity in response to P-site codon-anticodon mismatches"

**Supplementary data file contains**

**Figures S1-4**

**Tables S1, S2**

**Data deposition:** X-ray crystallography, atomic coordinates, and structure factors have been deposited in the Protein Data Bank, www.pdb.org (PDB codes 8FOM, 8FON)

**Key words:** ribosome, miscoding, near cognate, mRNA, tRNA, fidelity, translation

### **Supplementary Figures**

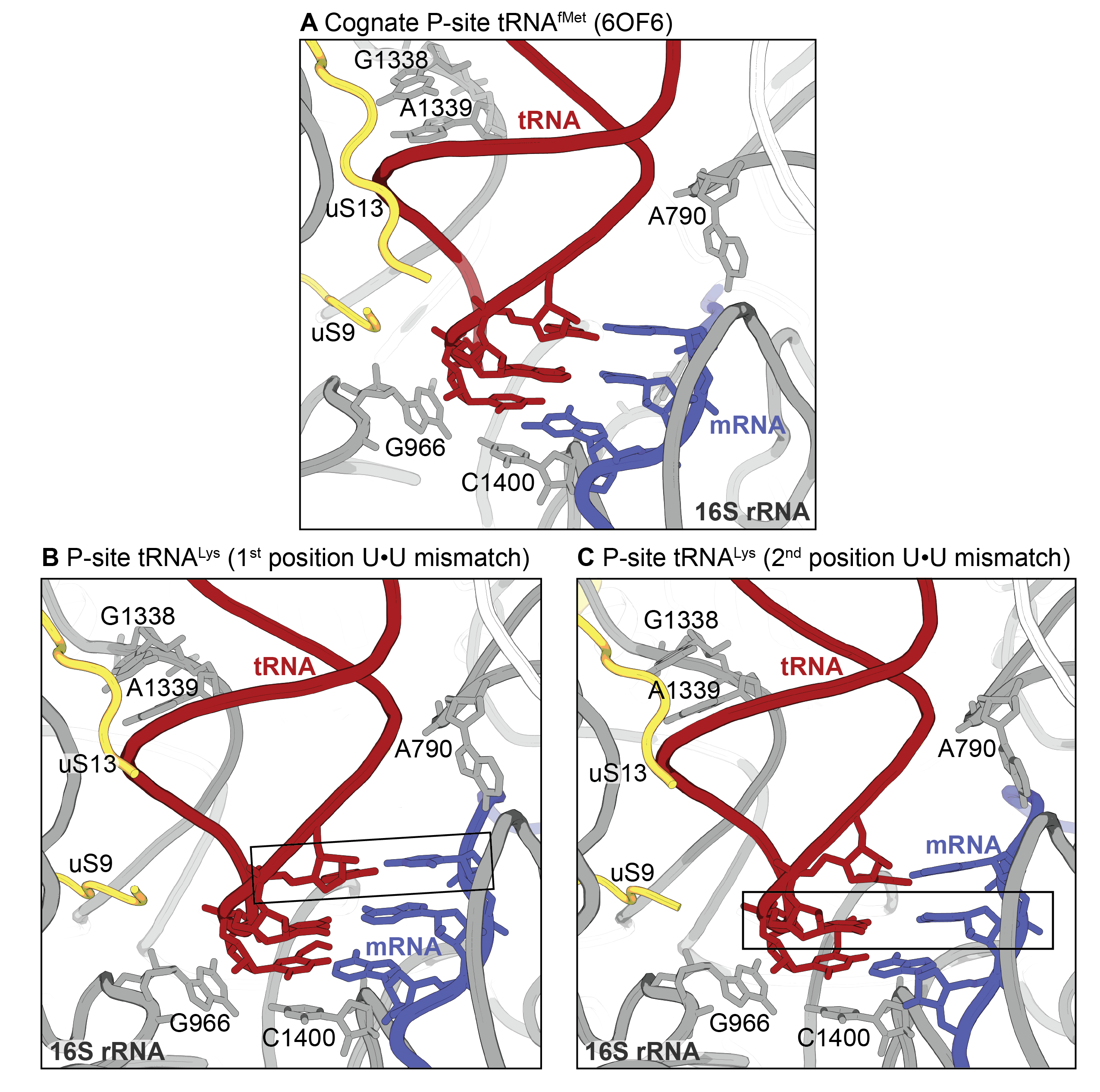

**Figure S1. The ribosomal P site does not recognize the mismatches in the codon-anticodon interaction.** The ribosomal environment of a cognate P-site tRNA^fMet^ bound to the start codon in the P site (PDB code 6OF6) (A) is similar to the two structures containing P-site mismatches at the first (B) or second position (C). The three base pairs of the codon-anticodon interaction are not extensively probed in the P site, and the P site is not known to have proofreading capability like that of the A site. In all three structures shown here, the P-site rRNA nucleotides and ribosomal proteins (r-proteins) critical for tRNA binding and translocation do not have any structural response to the mismatch in the first (B) and second position (C) of the codon-anticodon interaction: 16S rRNA nucleotides G966 and C1400 pack against the third anticodon-codon interaction similarly in all three structures, A790 is in the same conformation, G1338 and A1339 grips the stem of the anticodon loop, and r-proteins uS9 and uS13 tails form the tRNA binding site in the same way as when a correct tRNA is bound (A).

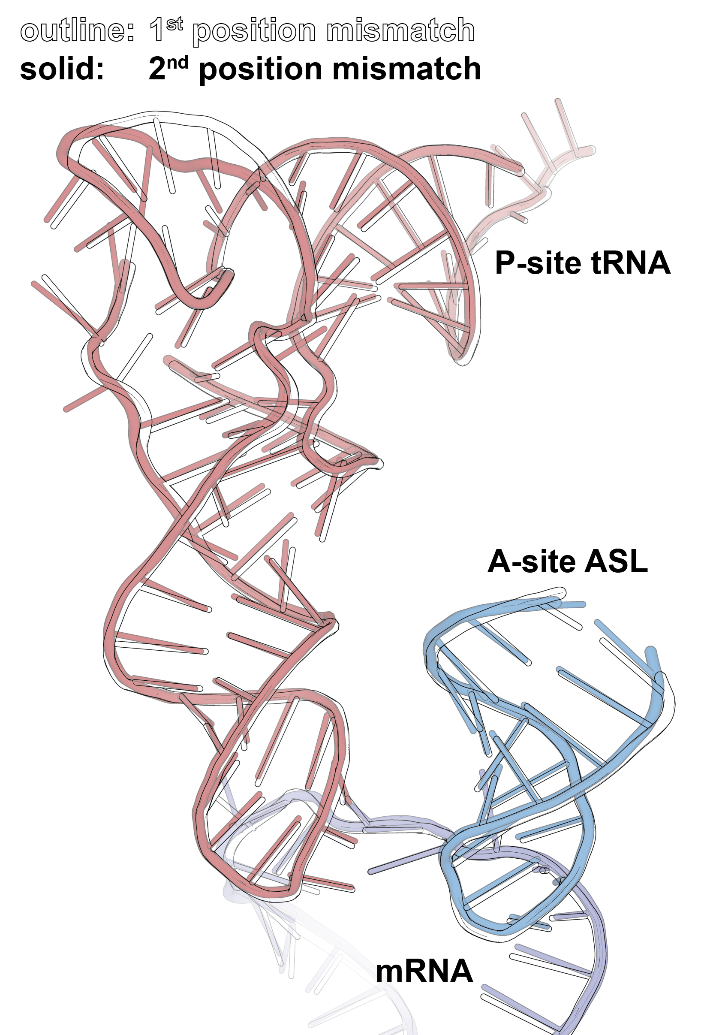

**Figure S2**. **Overlay of the P-site tRNAs of two structures in this study shows similar tRNA conformations**. The tRNA^Lys^ bound to the 1^st^ position mismatch (in black outline) looks almost identical to when it is bound to the 2^nd^ position mismatch (in red). The A-site ASL and the mRNA are also similar, with the overall all-atom root-mean-square deviation (RMSD) of the P-site tRNA, A-site ASL, and the mRNA being 0.357 Å.

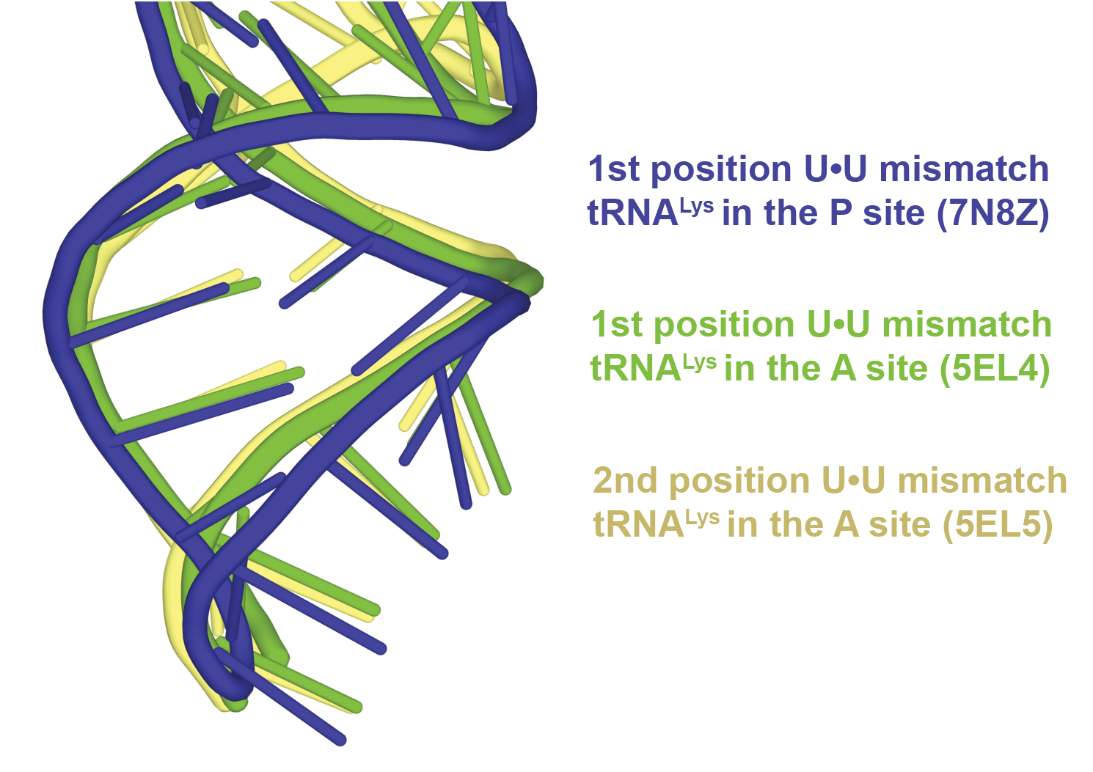

**Figure S3. Comparison of tRNA^Lys^-mRNA codon mismatches.** The anticodon stem loop conformation of the P-site tRNA^Lys^ 1^st^ position mismatch is similar as compared to the A-site tRNA^Lys^ with a 1^st^ (green) and 2^nd^ (yellow) position U•U mismatches (Rozov et al. 2016).

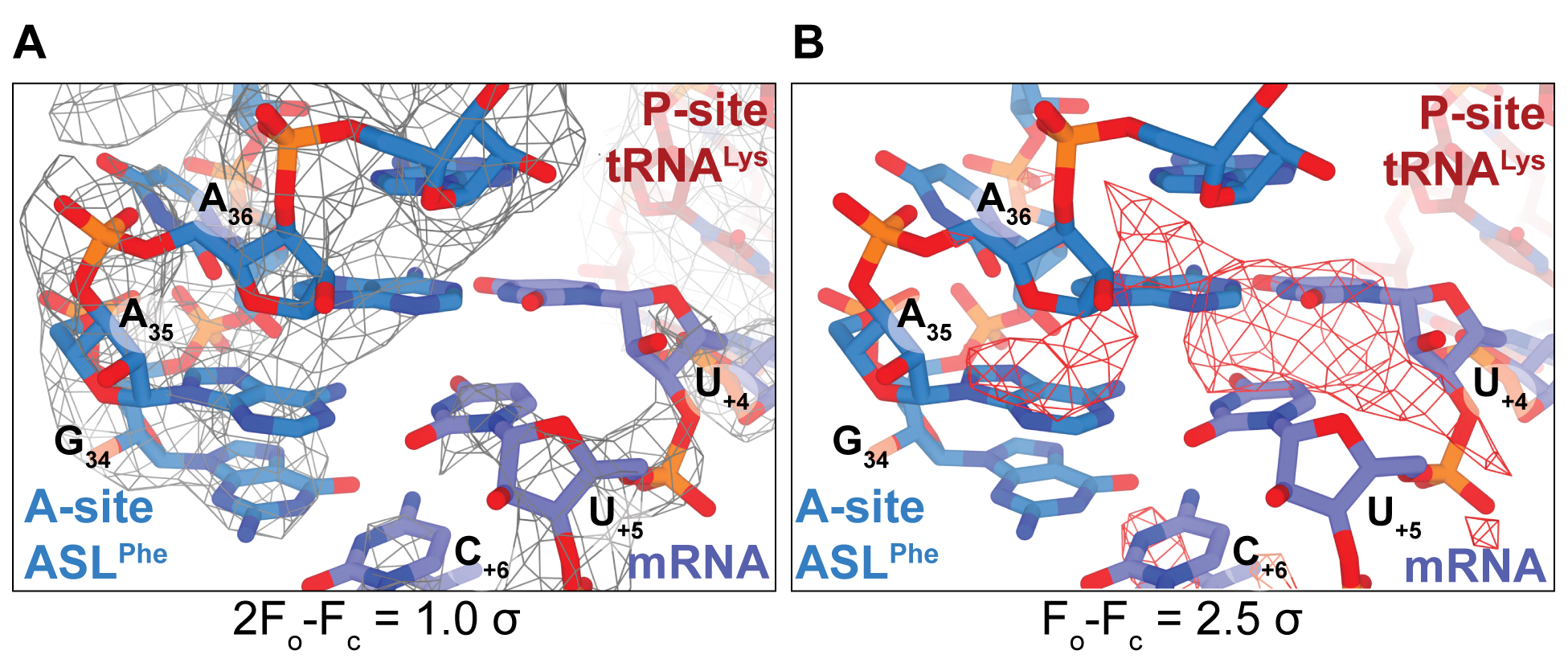

**Figure S4. Electron density maps after refinement of the structure containing the 2^nd^ position P-site mismatch when U_+4_ and A_36_ are modeled to form a base pair in the A site.** A. The gray mesh overlay represents the 2F_o_-F_c_ map contoured at 1.0 σ showing lack of density supporting the U_+4_ and A_36_ forming a Watson-Crick base pair. B. The red mesh overlay represents the F_o_-F_c_ map contoured at 2.5 σ showing strong negative difference density for the modeled base pair.

**Table S1 | Data collection and refinement statistics**

|  | **P-site tRNA^Lys^  UAA codon** | **P-site tRNA^Lys^  AUA codon** |
| --- | --- | --- |
| PDB code | 8FOM | 8FON |
| **Data collection** |  |  |
| Wavelength (Å) | 0.97890 | 0.97890 |
| Space group | P2_1_2_1_2_1_ | P2_1_2_1_2_1_ |
| Cell dimensions |  |  |
| *a, b, c (Å)* | 209.7, 449.5, 617.3 | 210.6. 447.2, 618.1 |
| *α, β, γ (°)* | 90, 90, 90 | 90, 90, 90 |
| Resolution (Å) | 137.3-3.58 (3.708-3.58) | 173.5-3.64 (3.77-3.64) |
| R_merge_ | 0.15 (0.86) | 0.19 (1.07) |
| R_pim_ | 0.094 (0.52) | 0.096 (0.54) |
| *I/σI* | 8.66 (1.70) | 7.28 (1.55) |
| CC_1/2_ | 0.997 (0.448) | 0.997 (0.438) |
| Completeness (%) | 98.3 (99.0) | 95.7 (97.5) |
| Redundancy | 3.4 (3.5) | 4.4 (4.4) |
| **Refinement** |  |  |
| Total reflections | 2,270,192 | 2,696,566 |
| Reflections used in refinement | 667,719 | 618,599 |
| *R*_work_/*R*_free_ (%) | 20.6/25.4 | 23.1/27.9 |
| No. atoms |  |  |
| Macromolecules | 291,264 | 291,330 |
| Ligands | 922 | 1,267 |
| B-factors |  |  |
| Macromolecules | 119.8 | 124.3 |
| Ligands | 61.1 | 50.6 |
| Clashscore | 12.7 | 5.5 |
| R.m.s. deviations |  |  |
| Bond lengths (Å) | 0.005 | 0.004 |
| Bond angles (°) | 0.93 | 0.87 |
| Ramachandran plot (%) |  |  |
| Favored regions (%) | 91.2 | 92.6 |
| Allowed regions (%) | 7.60 | 6.87 |
| Outliers (%) | 1.18 | 0.51 |

*Values in parentheses are for highest-resolution shell.

**Table S2 | RNA sequences used in this study.**

| mRNA P-site UAA codon | 5’- GGC AAG GAG GUA GGG AUG **UAA** UUC AAA -3’ |
| --- | --- |
| mRNA P-site AUA codon | 5’- GGC AAG GAG GUA GGG AUG **AUA** UUC AAA -3’ |
| ASL^Phe^ (GAA anticodon) | 5’- GGG GAU U**GA A**AA UCC CC -3' |
